# In-depth blood proteome profiling analysis revealed distinct functional characteristics of plasma proteins between severe and non-severe COVID-19 patients

**DOI:** 10.1101/2020.08.18.255315

**Authors:** Joonho Park, Hyeyoon Kim, So Yeon Kim, Yeonjae Kim, Jee-Soo Lee, Moon-Woo Seong, Dohyun Han

## Abstract

The severe acute respiratory syndrome coronavirus 2 (SARS-CoV-2) has infected over ten million patients worldwide. Although most coronavirus disease 2019 (COVID-19) patients have a good prognosis, some develop severe illness. Markers that define disease severity or predict clinical outcome need to be urgently developed as the mortality rate in critical cases is approximately 61.5%. In the present study, we performed indepth proteome profiling of undepleted plasma from eight COVID-19 patients. Quantitative proteomic analysis using the BoxCar method revealed that 91 out of 1,222 quantified proteins were differentially expressed depending on the severity of COVID-19. Importantly, we found 76 proteins, previously not reported, which could be novel prognostic biomarker candidates. Our plasma proteome signatures captured the host response to SARS-CoV-2 infection, thereby highlighting the role of neutrophil activation, complement activation, platelet function, and T cell suppression as well as proinflammatory factors upstream and downstream of interleukin-6, interleukin-1B, and tumor necrosis factor. Consequently, this study supports the development of blood biomarkers and potential therapeutic targets to aid clinical decision-making and subsequently improve prognosis of COVID-19.

## INTRODUCTION

Coronavirus disease 2019 (COVID-19) pandemic is an unprecedented global health threat caused by severe acute respiratory syndrome coronavirus 2 (SARS-CoV-2). Recent studies have reported an astonishing case fatality rate of 61.5% for critical cases, increasing sharply with age and in patients with underlying comorbidities [1]. The severity and increasing number of cases, the medical services face immense pressure, and there is a shortage of intensive care resources. To curtail the pandemic and return to normalcy, it is essential to find markers that define the disease severity, have prognostic value, or predict a specific phase of the disease. Unfortunately, no prognostic biomarkers are presently available that can distinguish patients requiring immediate medical attention and estimate their associated mortality rates. Nevertheless, Yan et al. [2] reported that blood-borne marker panels can identify the mortality rate in individual patients more than 10 days in advance with >90% accuracy. Moreover, they suggested that tissue damage markers can be leveraged to predict COVID-19 outcomes.

Mass spectrometry (MS)-based proteomics may potentially be used as an ideal technology in this situation as it can quickly deliver substantial amounts of clinical and biological information from blood plasma or serum in an untargeted manner [3,4]. Furthermore, these MS-based proteomic workflows for biomarker discovery and profiling are well established [3]. However, only two studies have presently applied proteomics to the serum of COVID-19 patients with moderate proteome depth [5,6]. Therefore, detailed understanding of the in-depth proteome of plasma or serum is necessary to develop prognostic or predictive protein markers.

In the present study, we performed in-depth proteome profiling of undepleted plasma samples using the BoxCar acquisition method [7] from an exploratory cohort comprising 8 COVID-19 patients to identify candidate biomarkers for evaluating the disease severity.

## RESULTS AND DISCUSSION

### 2.1. Label-free quantification of plasma samples

Here, we report the in-depth plasma proteome data of the Korean COVID-19 cohort. Our dataset was generated using the plasma samples collected from eight COVID-19 positively confirmed patients, including three non-severe (mild) and five severe cases (Supplementary Table S1). The plasma proteome was analyzed via ultra-high-resolution LC-MS and the proteins whose expression revealed significant differences were discovered (Figure 1). We provided this in-depth proteome as a cornerstone to the communities doing research on COVID-19.

**Figure 1.**
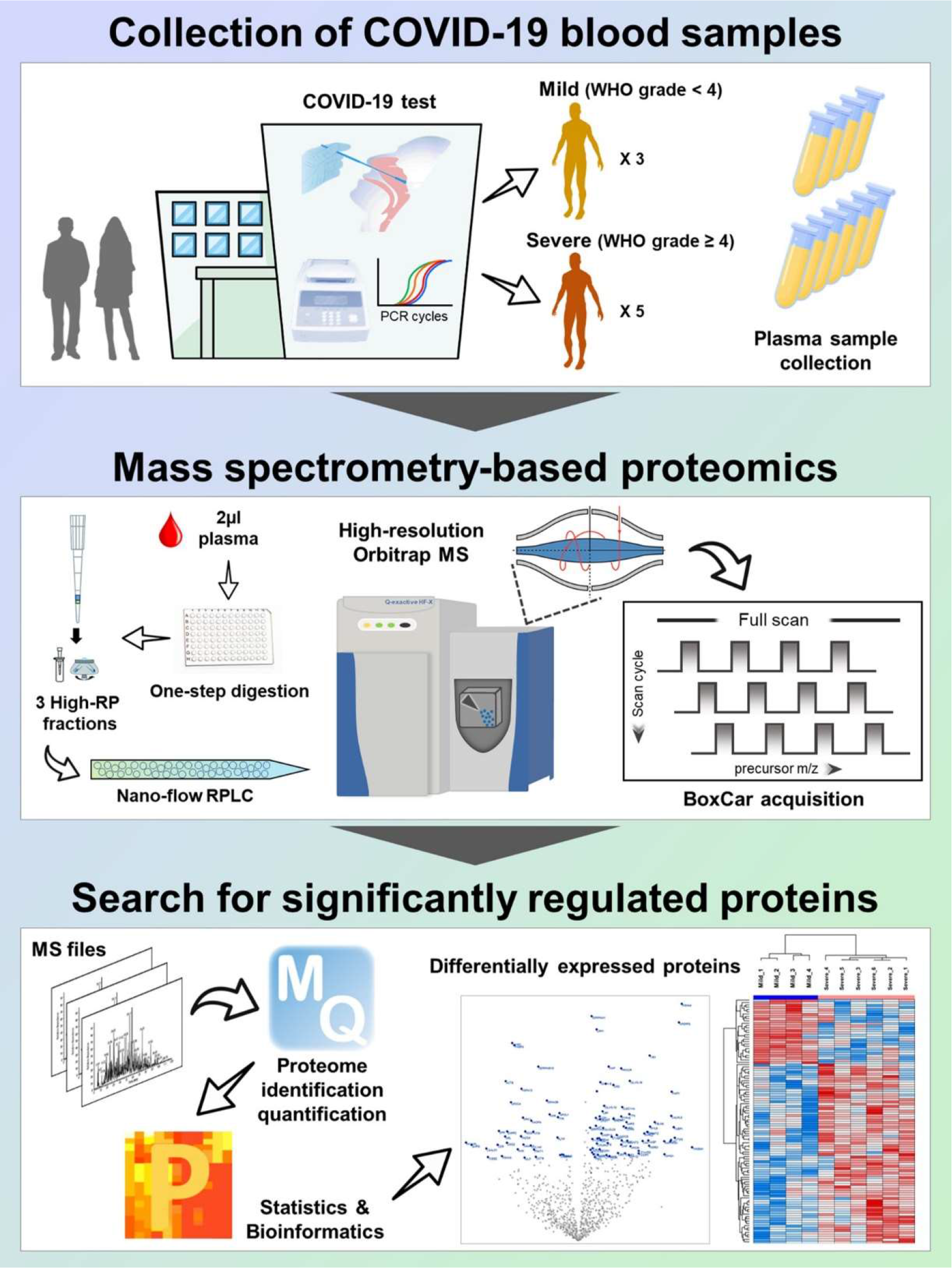
Overall scheme of in-depth plasma proteome profiling

To increase the proteome depth, we performed label-free quantification based on BoxCar acquisition using a small amount (2 μl) of plasma sample without high-abundant protein depletion. In total, 1,639 proteins were identified at the protein FDR 1% level. An average of 1,064 proteins were quantified in the individual samples (Figure 2a). The details of the identified and quantified proteins are listed in Supplementary Table S2. To identify the differences within and between groups, the protein profiles were plotted as multi-scatter plots, and the Pearson correlation coefficient (PCC) value between proteome pairs was calculated (Supplementary Figure S1). The intra-group correlation displayed average PCCs of 0.82 and 0.80 in mild and severe groups, respectively. The average PCC value of inter-group between the mild and severe was 0.78. Presumably, the differences between groups were slightly larger than those within groups. The overall mass spectrometric intensities presented no significant differences between all samples, although the plasma samples were prepared based on equal volume and not on the amount of plasma protein (Supplementary Figure S2). Interestingly, principal component analysis (PCA) performed using all identified plasma proteomes presented clear separation of the samples, indicating that the plasma protein expression was considerably altered based on the clinical symptoms (Figure 2b)

**Figure 2.**
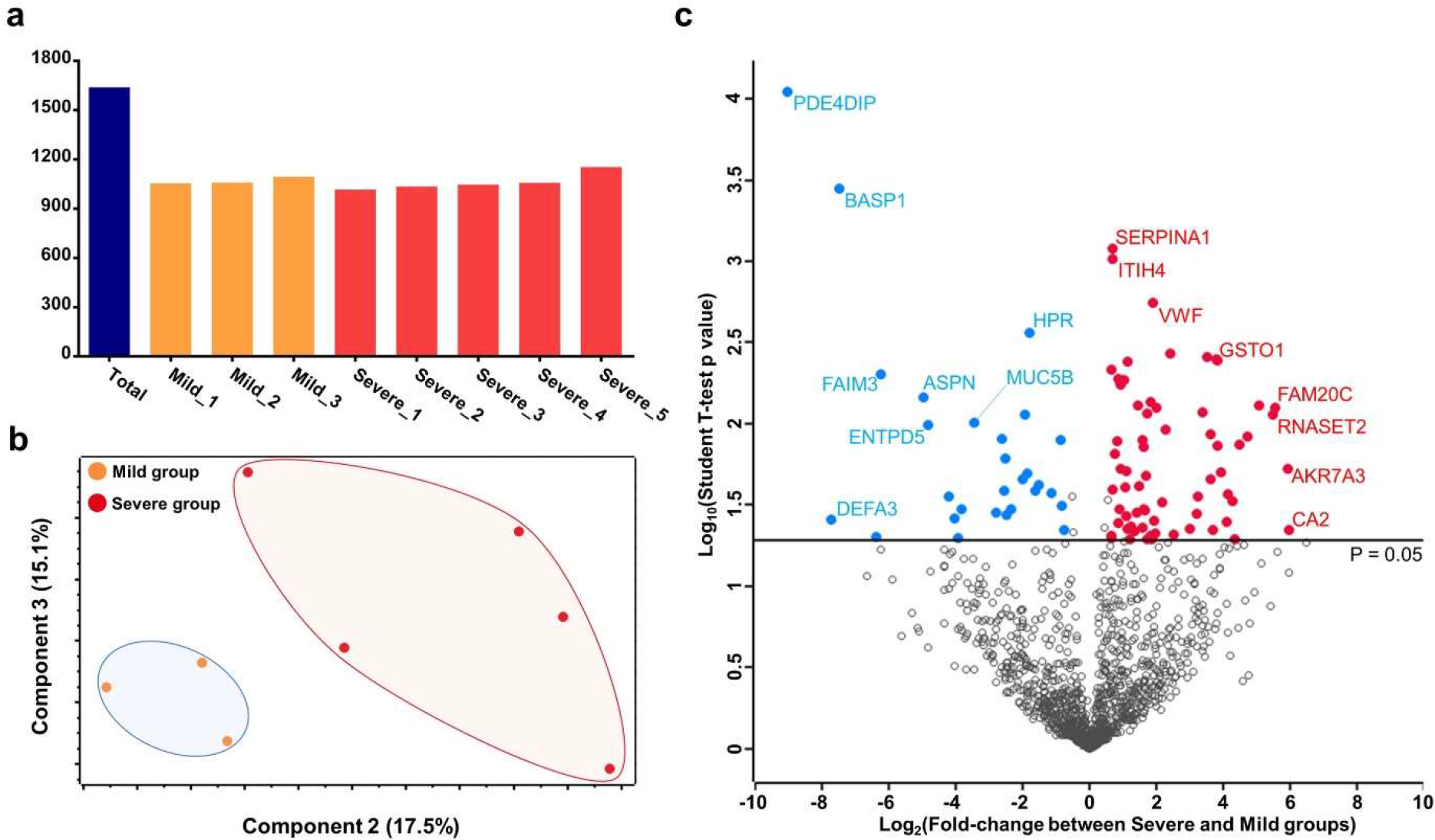
Plasma proteome data generated using COVID-19 infected patients. (a) The number of proteins identified in each plasma sample as well as the number of total identifications is plotted. (b) The result of principle component analysis (PCA) is depicted as 2-dimensional diagram. (c) The log-transformed student’s t-test p-value of each protein is plotted against the log-transformed fold-change. The middle line indicates the p-value cut-off, 0.05. The proteins with high significance (top eight) were labeled.

After considering the proteins quantified by at least 50% in either the mild or severe COVID-19 groups, 1,222 proteins were subjected to statistical analysis. Statistical tests revealed that the expression of 91 proteins significantly differed between mild and severe groups (Student’s t-test, p-value < 0.05, and |fold-change| > 1.5) (Figure 2c). These 91 proteins were regarded as DEPs and are summarized in Supplementary Table S3.

### 2.2 Comparison with previous studies

After the outbreak of COVID-19, two articles that intensively explored proteomics to discover blood biomarkers for COVID-19 have been published. Shen et al. analyzed the serum proteome and metabolome in a Chinese cohort (N = 118), suggesting a set of proteins as serum biomarkers for classifying COVID-19 patients [6]. Moreover, Messner et al. developed a high-throughput proteome analysis method and reported numerous significant proteins that distinguished the severe group from the non-severe group [5]. To verify the comprehensiveness of the proteome, our protein identifications and DEPs were compared with those of the previously published articles. The comparative analysis revealed that our proteome data covered most of the previous datasets, overlapping 71% and 72% of the proteome from Messner et al.’s dataset and Shen et al.’s dataset, respectively (Figure 3). Furthermore, we found 76 significant proteins, previously not reported, which could be novel biomarker candidates; however, the results differed from each other. Interestingly, although proteins such as C-reactive protein (CRP), serum amyloid A-1 (SAA1), protein Z-dependent protease inhibitor (SERPINA10), and albumin (ALB) were previously reported as promising marker candidates, these proteins could not fit into our criteria for differential expression. Presumably, the temporal gap between the blood sample collection time and the first symptom may be a reason for this disparity. The blood samples in our cohort were collected approximately 3 weeks after the first symptom, and the patients were treated with medications during this period. Therefore, the severe symptoms of the patients would have been alleviated, and thus the level of these proteins associated with acute responses would be restored to mild-symptom patients. Although numerous proteins were excluded, the expression of CRP, SAA1, Complement factor B (CFB), Cofilin-1 (CFL1), Complement C2 (C2), Leucine-rich alpha-2-glycoprotein (LRG1), Apolipoprotein C-I (APOC1), and Serotransferrin (TF) revealed expression trend consistent with those of other studies with medium significance (p-value < 0.1) (Supplementary Figure S3). The significant proteins reported in the two published articles are listed in Supplementary Table S4.

**Figure 3.**
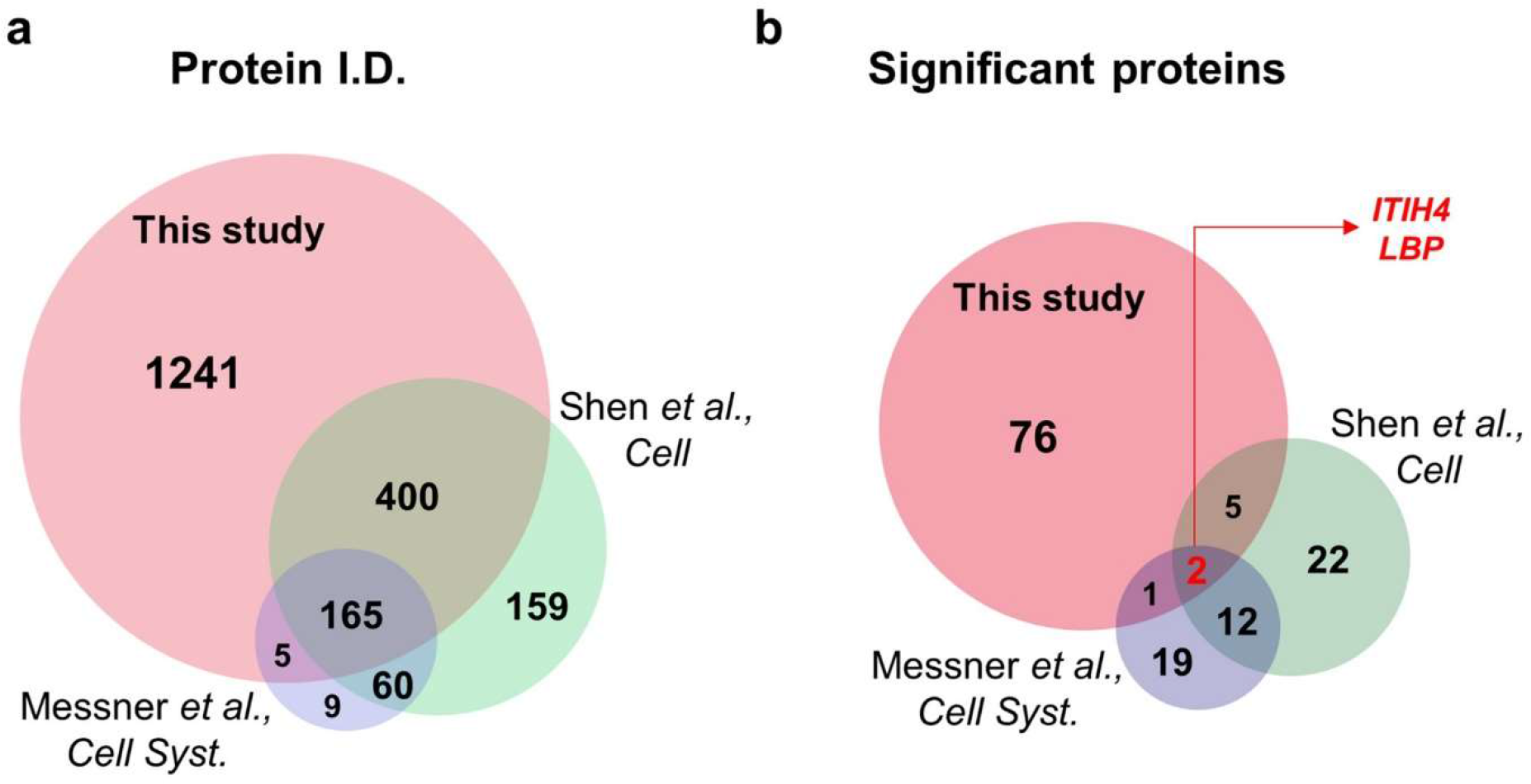
Comparison with other SARS-CoV-2 blood proteome datasets. (a) The list of protein identification in our data is compared to that of previously published papers (Shen et al., 2020, Cell and Messner et al., 2020, Cell systems). (b) The significant proteins proposed in each dataset were compared. Two proteins Inter-alpha-trypsin inhibitor heavy chain H4 (ITIH4) and Lipopolysaccharide-binding protein (LBP) were commonly reported as significant.

### 2.3 Functional characteristics distinguish the severe COVID-19 group from the mild group

To recapitulate the functional characteristics of differentially expressed plasma proteome and further suggest potential therapeutic targets for COVID-19, we investigated the biological functions and signaling pathways associated with DEPs. The over-representative analysis using EnrichR [8] based on bioPlanet and Wikipathway database derived numerous biological functions and signaling pathways satisfying the statistical criteria (Fisher’s exact test p-value < 0.05) (Supplementary Table S5). Notably, the functions related to the neutrophils and blood coagulation were mostly significant (Figure 4a). For example, the function “neutrophil activation involved in immune response” was associated with nine upregulated proteins, including Polymeric immunoglobulin receptor (PIGR), Fructose-bisphosphate aldolase C (ALDOC), Heat shock cognate 71 kDa protein (HSPA8), Vesicle-associated membrane protein-associated protein A (VAPA), Ras GTPase-activating-like protein (IQGAP2), Serpin B10 (SERPINB10), Alpha-1-antitrypsin (SERPINA1), and Alpha-1-antichymotrypsin (SERPINA3), and was deduced as one of the most important functions (p-value = 1.12E-05). Other similar terms such as “neutrophil degranulation” and “neutrophil mediated immunity” were also enriched (p-value = 1.04E-05 and p-value = 1.19E-05, respectively). Recently, the role of neutrophils in severe COVID-19 has received immense attention. Specifically, a microarray-based study of SARS-CoV-2 infected a peripheral blood mononuclear cell (PMBC) and single cell analysis of epithelial and immune cells in COVID-19 patients revealed that the neutrophil markers were overexpressed, suggesting that the patients were under neutrophilia [9,10]. Furthermore, a meta-analysis based on gene network constructed from the published datasets derived several neutrophil-enriched genes [11]. Other articles have reported that neutrophils and their extracellular traps (Neutrophil extracellular traps, NETs) trigger COVID-19 [12,13]. Barnes et al. [12] argued that the NETs formed by expelled proteins and DNA by neutrophils play a crucial role in protecting the host; however, the excessive persistence of NETs induce a hyperinflammatory response and thus may damage the organs. Moreover, they suggested that NETs also contribute to cytokine storm by stimulating macrophages to secrete cytokines such as interleukin-1-beta (IL-1B) and interleukin-6 (IL-6). Based on this perspective, the treatment strategy for the regulation of NET in severe COVID-19 patients is deemed important.

**Figure 4.**
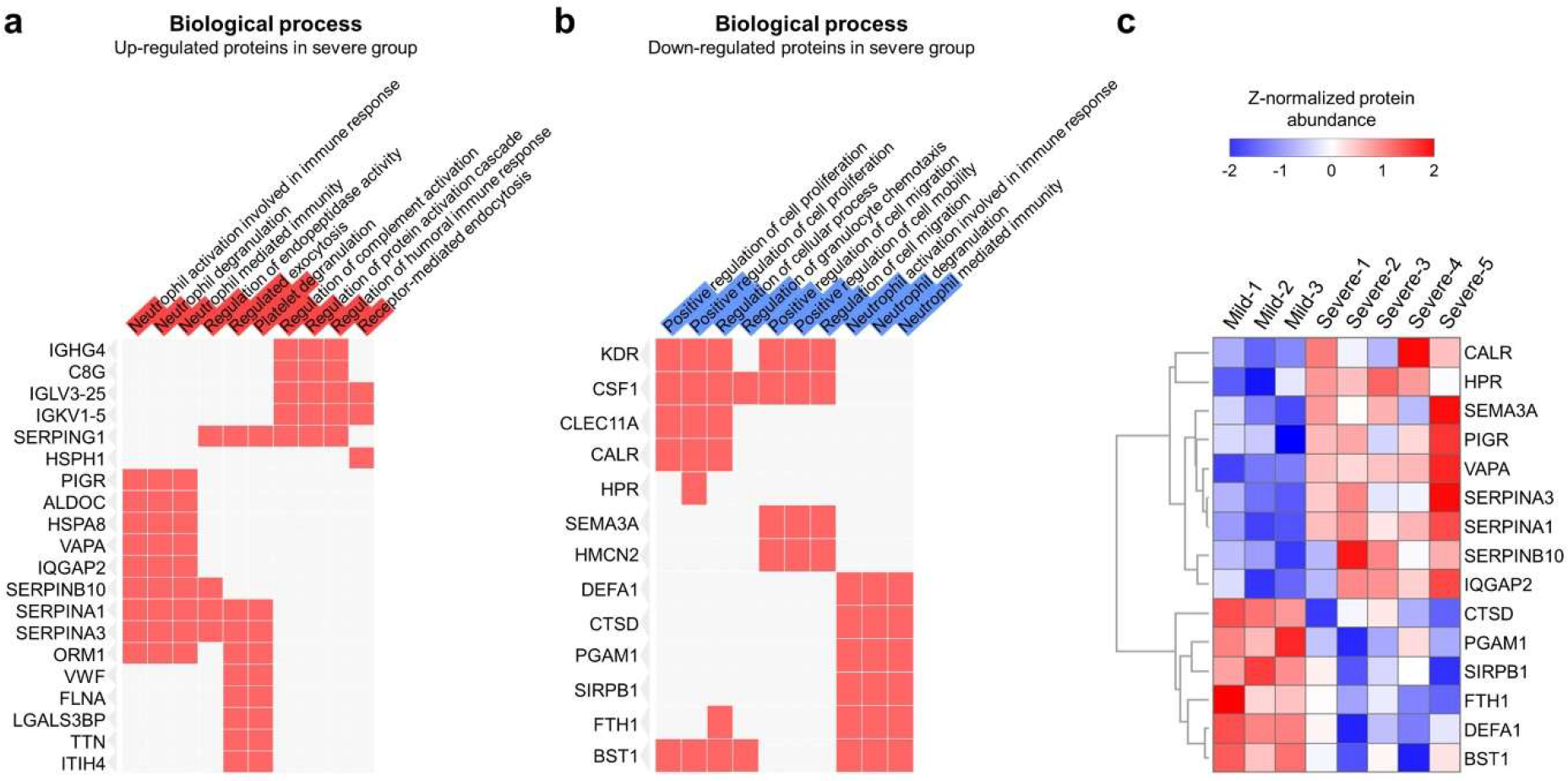
Functional analysis of differentially expressed proteins. (a, b) Biological function enrichment analysis was performed using the upregulated or downregulated DEPs. If the proteins on the left side were associated with the corresponding function, then they were filled with red. The level of significance of each biological function is displayed as the length of red (a) and blue (b) bars overlapped in the function term cell. (c) The expression levels of the 15 proteins associated with neutrophils were plotted.

Interestingly, identical functions were enriched in the downregulated proteins; Neutrophil defensin 3 (DEFA1), Cathepsin D (CTSD), Phosphoglycerate mutase 1 (PGAM1), Signal-regulatory protein beta-1 (SIRPB1), Ferritin heavy chain (FTH1), and ADP-ribosyl cyclase/cyclic ADP-ribose hydrolase 2 (BST1) enriched the “neutrophil activation involved in immune response” (p-value = 3.70E-05), “neutrophil degranulation” (p-value = 3.53E-05), and “neutrophil mediated immunity” (p-value = 3.87E-05) (Figure 4b). Our results revealed that the two protein groups are conflictingly affecting the activation of neutrophils (Figure 4c). The exact mechanism of these proteins in the SARS-CoV-2 infected blood cells need to be examined; however, these protein levels in plasma efficiently differentiate between the COVID-19 mild and severe groups, and thus, they could be suggested as potential prognostic markers.

Next, according to the study by Zhou et al., the proportion of coagulopathy increased significantly in the COVID-19 death group [14]. A meta-analysis report revealed that COVID-19-confirmed patients presented coagulopathy different from typical acute disseminated intravascular coagulopathy with the reduced levels of the fibrinogen and relatively minimal changes in platelet counts [15]. According to the report, significantly elevated D-dimer and fibrinogen levels were the most common finding in COVID-19-related coagulopathy [13,15]. Other studies have also reported similar phenomenon revealing that severely affected COVID-19 patients are under a hypercoagulable state [16–18]. However, platelet counts in COVID-19 patients are variable depending on the reported studies [15]. In the present study, upregulation of the proteins involved in regulating complement and coagulation activation and platelet degranulation in the severe group can be considered as a reflection of coagulopathy. Potentially, SARS-CoV-2 entry from angiotensin-converting enzyme-2 (ACE-2) can release intracellular angiotensin II, triggering platelet degranulation [19]. This resulted in inflammation and loss of platelets via deposition in peripheral microvascular beds, thereby heralding thrombocytopenia and intravascular coagulopathy in COVID-19 [20]. Presently, some experts cautiously suggest that severe COVID-19 patients require medications with anticoagulation drugs as adjunctive therapy to reduce severity and mortality [21], and our results support this treatment strategy.

Additionally, signaling pathway enrichment using EnrichR software also revealed that the upregulated proteins in the severe group were predicted to be involved mainly in pathways related to platelet function and coagulation, thereby further highlighting the importance of coagulation in COVID-19, as established by previous observations in patients with severe symptoms [13,15–17]. Furthermore, the metabolism pathway was presumed to be activated (Supplementary Figure S4a). In contrast, various signaling pathways were enriched in downregulated proteins, such as interleukin-2 (IL-2) signaling, signaling events mediated by T cell protein tyrosine phosphatase, and lysosome (Supplementary Figure S4b). Notably, a recent study reported that the inhibition of IL-2 signaling may relate to the decreased CD8+ T cells in critical COVID-19 patients [22]. Here, we observed downregulation of five proteins included in the IL-2 signaling pathway, such as Macrophage colony-stimulating factor 1 (CSF1), Vascular endothelial growth factor receptor 2 (KDR), Myomegalin (PDE4DIP), Fas apoptotic inhibitory molecule 3 (FAIM3), and CTSD, in critical patients, which may represent another potential therapeutic target for COVID-19.

To identify potential therapeutic targets, we used the molecule–molecule interaction causality information present in IPA. We constructed a regulator-target network using upstream regulators predicted to regulate the DEPs found in this study (Figure 5). As a result, eight upstream regulators that regulate 28 DEPs were discovered; of these, six regulators were increased and remaining two regulators were decreased. One of the most significant regulators is IL-6, and the immune response triggered by it is activated in severe COVID-19 patients, as reported in previous publications [5,6,23,24]. In addition to IL-6, IL-1B and tumor necrosis factor (TNF) are the major components of the cytokine storm commonly observed in severe COVID-19 patients [25]. The aforementioned results might be due to immune responses that naturally occur in COVID-19. Notably, proto-oncoproteins such as Cellular tumor antigen p53 (TP53) and the Myc proto-oncogene protein (MYC) family were predicted to regulate seven plasma DEPs. In particular, decrease in TP53 as well as increase in MYC, were predicted. Presumably, the regulation of these proteins may contribute to cell proliferation, particularly macrophages and neutrophils, in response to SARS-Cov-2 infection. In contrast, the regulated levels of DEPs, including Proprotein convertase 1 inhibitor (PCSK1N), Galectin-3-binding protein (LGALS3BP), Insulin-like growth factor-binding protein 3 (IGFBP3), and ADP-ribosyl cyclase/cyclic ADP-ribose hydrolase 2 (BST1), elicited a predicted increase in PDZ and LIM domain protein 2 (PDLIM2) levels. PDLIM2 has been reported to inhibit T cell development, which is in accordance with previous reports on reduced numbers and functional diversity of T cells in severe COVID-19 patients [26–28]. Moreover, the level of T cell receptor (TCR) complex was reduced in our analysis. Therefore, it can be considered that the quantity or function of T cells decreased as COVID-19 exacerbated. Nevertheless, our suggestion is based on in silico analysis; hence, the role of regulators and their target proteins in severe COVID-19 should be verified via further functional studies.

**Figure 5.**
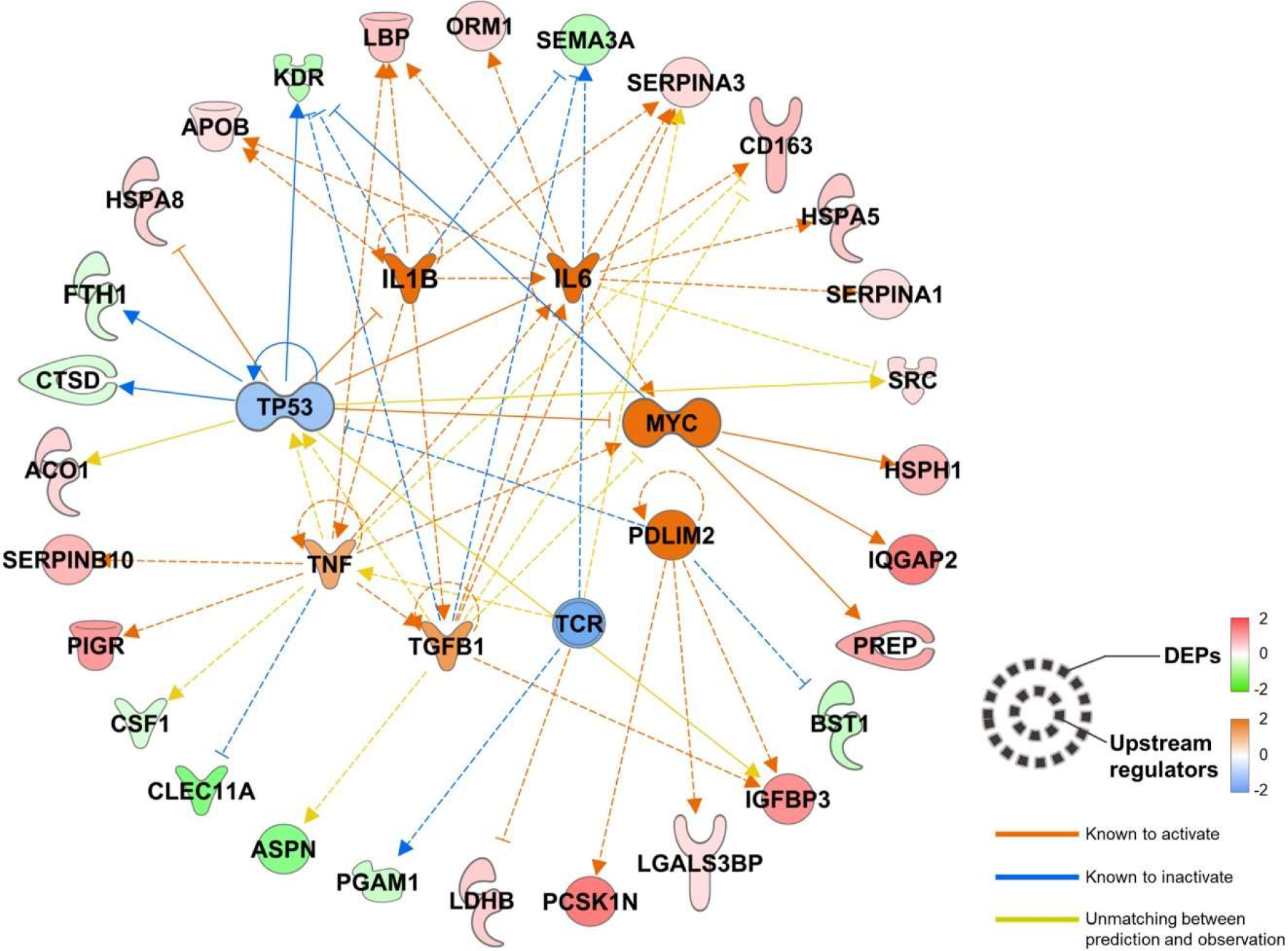
Upstream regulator–downstream plasma DEP network. The upstream regulators that were known or reported to regulate the input DEPs at the upper level were searched, and their status was calculated using the Ingenuity Pathway Analysis bioinformatics tool. Eight regulators were predicted to be upregulated and one to be downregulated. The upstream regulators, their regulation targets, that is, plasma DEPs in our dataset, and the functional connections between them were plotted.

## METHODS

### Patients and samples

Our team procured plasma samples from eight COVID-19 patients who visited the National Medical Center between February and April 2020 (Supplementary Table S1). All patients with positive results for SARS-CoV-2 infection, detected by analyzing the respiratory specimens via PCR, and willing to provide written informed consent were eligible for participation. Treatment and medical interventions follow the standard of care as recommended by the current international and Korean guidelines for COVID-19. The severity of COVID-19 in this study was classified based on the requirement of oxygen support (mild = no oxygen support; severe = oxygen support). The Ethics Committee and Institutional Review Board of the National Medical Center (IRB no. H-2002-111-002) approved all experimental procedures. All experiments were performed following the approved guidelines.

### Sample preparation

The protein digestion process was optimized to 2 μl of each plasma or serum sample as previously described with some modifications [29]. Briefly, 23 μl of digestion buffer, including reduction and alkylation reagents, was added to 2 μl of blood plasma or serum. The mixture was boiled for 25 min at 60 °C to denature and alkylate the proteins. After cooling to room temperature, proteins were digested at 37 °C overnight using a trypsin/LysC mixture at 100:1 protein-to-protease ratio. The second digestion was performed at 37 °C for 2 h using trypsin (enzyme-to-substrate ratio [w/w] of 1:1,000). All resulting peptides were acidified with 10% trifluoroacetic acid (TFA). The acidified peptides were loaded onto a homemade styrenedivinylbenzene reversed-phase sulfonate (SDB-RPS)-StageTips according to the previously described procedures [29,30]. The peptides were initially washed with 0.2% TFA, and were then eluted with 80% acetonitrile (ACN) containing 1% ammonia. The eluate was vacuum-centrifuged to dryness and stored at −80 °C.

### LC-MS/MS analysis

All LC-MS/MS analyses were performed using Quadrupole Orbitrap mass spectrometers, Q-exactive HF-X (Thermo Fisher Scientific, Waltham, MA, USA) coupled to an Ultimate 3000 RSLC system (Dionex, Sunnyvale, CA, USA) via a nanoelectrospray source, as described previously with some modifications [29,31]. Peptide samples were separated on a 2-column setup with a trap column (75 μm I.D. × 2 cm, C18 3 μm, 100 Å) and an analytical column (50 μm I.D. × 15 cm, C18 1.9 um, 100 Å). Prior to sample injection, the dried peptide samples were predissolved in solvent A (2% acetonitrile and 0.1% formic acid). After loading the samples onto the Nano LC, a 90-min gradient from 8% to 26% of solvent B (100% acetonitrile and 0.1% formic acid) was applied to all samples. The spray voltage was 2.0 kV in the positive ion mode, and the temperature of the heated capillary was set to 320 °C. MaxQuant.Live version 1.2 was used to perform BoxCar acquisition [7,32]. The MS1 resolution was set to 120,000 at m/z 200 for BoxCar, and the acquisition cycle comprised two BoxCar scans at 12 boxes (scaled width, 1 Th overlap) with a maximum ion injection time of 20.8 per box with the individual AGC target set to 250,000.

### Spectral library generation

To generate spectral libraries for the BoxCar, 24 data-dependent acquisition (DDA) measurements of the immunodepleted plasma samples were performed. DDA spectra were searched using the Maxquant against Uniprot Human Database (December 2014, 88,657 entries) and the iRT standard peptide sequence.

### Data processing for label-free quantification

Mass spectra were processed using MaxQuant version 1.6.1.0 [33]. MS/MS spectra were searched against the Human Uniprot protein sequence database (December 2014, 88,657 entries) using the Andromeda search engine [34]. In the global parameter, the BoxCar was set as the experimental type. Primary searches were performed using a 6-ppm precursor ion tolerance for analyzing the total protein levels. The MS/MS ion tolerance was set at 20 ppm. Cysteine carbamide-methylation was set as a fixed modification. N-Acetylation of proteins and oxidation of methionine were set as variable modifications. Enzyme specificity was set to complete tryptic digestion. Peptides with a minimum length of six amino acids and up to two missed-cleavages were considered. The required false discovery rate (FDR) was set to 1% at the peptide, protein, and modification levels. To maximize the number of quantification events across samples, we enabled the “Match between Runs” option on the MaxQuant platform. The MS proteomics data have been deposited to the ProteomeXchange Consortium via the PRIDE [35] partner repository with the data set identifier PXD020354.

### Statistical analysis

For pair-wise analysis in plasma experiments, data were statistically analyzed using Perseus software [36]. Initially, proteins only identified by site, reverse, and contaminants were removed. The expression level of proteins in the plasma was estimated by determining their intensity based absolute quantification (iBAQ) values calculated using Maxquant software. After log2 transformation was conducted for these values, valid values were filtered using proteins with a minimum of 50% quantified values in at least one symptom group. Missing values of the proteins were imputed based on a normal distribution (width = 0.5, down-shift = 1.8) to simulate signals of low-abundance proteins. Two-sided t-tests were performed for pairwise comparisons of proteomes to detect differentially expressed proteins (DEPs) with significant filtering criteria (p-value < 0.05 and fold-change > 1.5). Protein abundances were subjected to z-normalization followed by hierarchical clustering with Pearson’s correlation distance.

### Bioinformatics analysis

Funtional gene ontology (GO) and pathway enrichment analysis were performed using the EnrichR online tool (http://amp.pharm.mssm.edu/Enrichr/) [8]. Upstream regulation and protein networks were evaluated via Ingenuity Pathway Analysis (IPA, QIAGEN, Hilden, Germany) based on the DEPs in the plasma experiment. The analytical algorithms embedded in IPA uses lists of DEPs to predict the biological processes and pathways. The statistical significance of both the gene ontology classification and enrichment analysis was determined by Fisher’s exact test. All statistical tests were two-sided, and p < 0.05 was considered as statistically significant.

## CONCLUSIONS

In conclusion, a class of emerging coronaviruses, including SARS-CoV-2, SARS, and MERS, caused worldwide health concerns. Although recent efforts can acquire the genetic sequence of the virus and initial data on the epidemiology and clinical consequences of SARS-CoV-2, numerous important questions remain unanswered, including its origin, extent, duration of transmission in humans, ability to infect other animal hosts, and the spectrum and pathogenesis of human infections. In particular, insufficient biochemical knowledge will make it difficult to identify the biomarkers and to define point-of-care clinical classifiers. MS-based proteomics can present valuable and unbiased information about disease progression and therapeutic targets, without prior knowledge about the etiologies and biomolecules.

To the best of our knowledge, with a total of 1,639 proteins identified and 1,222 proteins statistically analyzed, this is the first comprehensive study of the plasma proteome for COVID-19 patients, which provides a unique insight into the altered protein circulation based on the severity of COVID-19. We identified 91 differentially expressed plasma proteins between the mild and severe groups of COVID-19 and demonstrated the potential of plasma proteome signatures. The proteome signatures captured the host response to COVID-19 infection, highlighting the role of neutrophil activation, complement activation, and platelet function. Furthermore, our bioinformatics analysis indicated a high specificity of several inflammatory modulators, particularly IL-6, IL-1B, and TNF. Overrepresentation of the suppressing factor of T cells (PDLIM2) was also predicted. This study had certain limitation including the sample size of COVID-19 patients; the clinical significance and statistical power would increase with more patients. Nevertheless, our in-depth plasma proteome may provide insights into the development of prognostic biomarkers to support clinical decisionmaking as well as potential therapeutic targets.

## DATA AVAILABILITY

The mass spectrometry data generated during and/or analyzed during the current study are deposited and available in the PRIDE Archive (http://www.ebi.ac.uk/pride/archive) with the dataset identifier; PXD020354.

## ACKNOWLEDGMENTS

We glad to acknowledge the contribution of the research staffs as well as other collaborators affiliated to Seoul National University and Seoul National University Hospital. We also thank all the patients who participated in the study. This work was supported by the National Research Foundation grant (NRF-2019R1F1A1058753 and NRF-2020R1A5A1019023), funded by the Korean Government (MSIP). Finally, this work was supported by grant no 0420190860 from the SNUH Research Fund.

## AUTHOR CONTRIBUTIONS

D.H. and M-W.S. conceived, designed, and supervised the overall study. M-W.S., S.Y.K., Y.K. and J-S.L. contributed in recruiting cohorts and collecting the clinical samples. H.K., D.H., and J.P. collected the proteomics data and carried out the quantitative analyses. After J.P. and D.H. prepared the manuscript, all authors contributed in revising it. All authors approved the submission of the final manuscript and agreed to be responsible for all aspects of the work.

## ADDITIONAL INFORMATION

The authors who have taken part in this study declared that they do not have anything to disclose regarding funding or conflict of interest with respect to this manuscript.

## SUPPLEMENTARY INFORMATION

Supplementary Figure S1. Correlation plot in plasma experiment

Supplementary Figure S2. Profile plot of all proteins across the plasma samples

Supplementary Figure S3. The expression summary for the proteins reported as significant in previous papers

Supplementary Figure S4. Pathway enrichment using up- or down-regulated DEPs

Supplementary Table S1. Clinilcal information of pilot COVID-19 cohort

Supplementary Table S2. Total list of the identified proteins in plasma experiment

Supplementary Table S3. Differentially expressed proteins between mild and severe group

Supplementary Table S4. Comparison of proteome data with other studies

Supplementary Table S5. Gene ontology and pathway enrichment analysis using EnrichR

## REFERENCES

1 Yang, X. et al. Clinical course and outcomes of critically ill patients with SARS-CoV-2 pneumonia in Wuhan, China: a single-centered, retrospective, observational study. Lancet Respir Med 8, 475–481 (2020).

2 Yan, L. et al. An interpretable mortality prediction model for COVID-19 patients. Nature Machine Intelligence 2, 283–288 (2020).

3 Geyer, P E., Holdt, L. M., Teupser, D. & Mann, M. Revisiting biomarker discovery by plasma proteomics. Mol Syst Biol 13, 942 (2017).

4 Whetton, A. D., Preston, G. W., Abubeker, S. & Geifman, N. Proteomics and Informatics for Understanding Phases and Identifying Biomarkers in COVID-19 Disease. J Proteome Res (2020).

5 Messner, C. B. et al. Ultra-High-Throughput Clinical Proteomics Reveals Classifiers of COVID-19 Infection. Cell Syst 11, 11–24 e14 (2020).

6 Shen, B. et al. Proteomic and Metabolomic Characterization of COVID-19 Patient Sera. Cell 182, 59–72 e15 (2020).

7 Meier, F., Geyer, P. E., Virreira Winter, S., Cox, J. & Mann, M. BoxCar acquisition method enables single-shot proteomics at a depth of 10,000 proteins in 100 minutes. Nat Methods 15, 440–448 (2018).

8 Kuleshov, M. V. et al. Enrichr: a comprehensive gene set enrichment analysis web server 2016 update. Nucleic Acids Res 44, W90–97 (2016).

9 Chua, R. L. et al. COVID-19 severity correlates with airway epithelium-immune cell interactions identified by single-cell analysis. Nat Biotechnol (2020).

10 Hemmat, N. et al. Neutrophils, Crucial, or Harmful Immune Cells Involved in Coronavirus Infection: A Bioinformatics Study. Front Genet 11, 641 (2020).

11 Gardinassi, L. G., Souza, C. O. S., Sales-Campos, H. & Fonseca, S. G. Immune and Metabolic Signatures of COVID-19 Revealed by Transcriptomics Data Reuse. Front Immunol 11, 1636 (2020).

12 Barnes, B.J. et al. Targeting potential drivers of COVID-19: Neutrophil extracellular traps. J Exp Med 217 (2020).

13 Zuo, Y. et al. Neutrophil extracellular traps in COVID-19. JCI Insight 5 (2020).

14 Zhou, F. et al. Clinical course and risk factors for mortality of adult inpatients with COVID-19 in Wuhan, China: a retrospective cohort study. Lancet 395, 1054–1062 (2020).

15 Iba, T., Levy, J. H., Levi, M., Connors, J. M. & Thachil, J. Coagulopathy of Coronavirus Disease 2019. Crit Care Med (2020).

16 Iba, T. et al. The unique characteristics of COVID-19 coagulopathy. Crit Care 24, 360 (2020).

17 Marietta, M., Coluccio, V. & Luppi, M. COVID-19, coagulopathy and venous thromboembolism: more questions than answers. Intern Emerg Med (2020).

18 Thachil, J. et al. ISTH interim guidance on recognition and management of coagulopathy in COVID-19. J Thromb Haemost 18, 1023–1026 (2020).

19 Biancardi, V. C., Bomfim, G. F., Reis, W. L., Al-Gassimi, S. & Nunes, K. P. The interplay between Angiotensin II, TLR4 and hypertension. Pharmacol Res 120, 88–96 (2017).

20 Kuchi Bhotla, H. et al. Platelets to surrogate lung inflammation in COVID-19 patients. Med Hypotheses 143, 110098 (2020).

21 Tang, N. et al. Anticoagulant treatment is associated with decreased mortality in severe coronavirus disease 2019 patients with coagulopathy. J Thromb Haemost 18, 1094–1099 (2020).

22 Shi, H. et al. The inhibition of IL-2/IL-2R gives rise to CD8(+) T cell and lymphocyte decrease through JAK1-STAT5 in critical patients with COVID-19 pneumonia. Cell Death Dis 11, 429 (2020).

23 Aziz, M., Fatima, R. & Assaly, R. Elevated interleukin-6 and severe COVID-19: A metaanalysis. J Med Virol (2020).

24 Ulhaq, Z. S. & Soraya, G. V. Interleukin-6 as a potential biomarker of COVID-19 progression. Med Mal Infect 50, 382–383 (2020).

25 Del Valle, D.M. et al. An inflammatory cytokine signature helps predict COVID-19 severity and death. medRxiv (2020).

26 De Biasi, S. et al. Marked T cell activation, senescence, exhaustion and skewing towards TH17 in patients with COVID-19 pneumonia. Nat Commun 11, 3434 (2020).

27 Vabret, N. et al. Immunology of COVID-19: Current State of the Science. Immunity 52, 910–941 (2020).

28 Zheng, H.Y. et al. Elevated exhaustion levels and reduced functional diversity of T cells in peripheral blood may predict severe progression in COVID-19 patients. Cell Mol Immunol 17, 541–543 (2020).

29 Rhee, S. J. et al. Comparison of serum protein profiles between major depressive disorder and bipolar disorder. BMC Psychiatry 20, 145 (2020).

30 Kim, H. et al. An efficient method for high-pH peptide fractionation based on C18 StageTips for in-depth proteome profiling. Analytical Methods 11, 4693–4698 (2019).

31 Kim, Y.S. et al. In-Depth, Proteomic Analysis of Nasal Secretions from Patients With Chronic Rhinosinusitis and Nasal Polyps. Allergy Asthma Immunol Res 11, 691–708 (2019).

32 Wichmann, C. et al. MaxQuant.Live Enables Global Targeting of More Than 25,000 Peptides. Mol Cell Proteomics 18, 982–994 (2019).

33 Tyanova, S., Temu, T. & Cox, J. The MaxQuant computational platform for mass spectrometry-based shotgun proteomics. Nat Protoc 11, 2301–2319 (2016).

34 Cox, J. et al. Andromeda: a peptide search engine integrated into the MaxQuant environment. J Proteome Res 10, 1794–1805 (2011).

35 Perez-Riverol, Y. et al. The PRIDE database and related tools and resources in 2019: improving support for quantification data. Nucleic Acids Res 47, D442–D450 (2019).

36 Tyanova, S. et al. The Perseus computational platform for comprehensive analysis of (prote)omics data. Nat Methods 13, 731–740 (2016).

