## Supplementary figures & tables for "In-depth blood proteome profiling analysis revealed distinct functional characteristics of plasma proteins between severe and non-severe COVID-19 patients": Supplementary Material_Figure S1-4.pdf

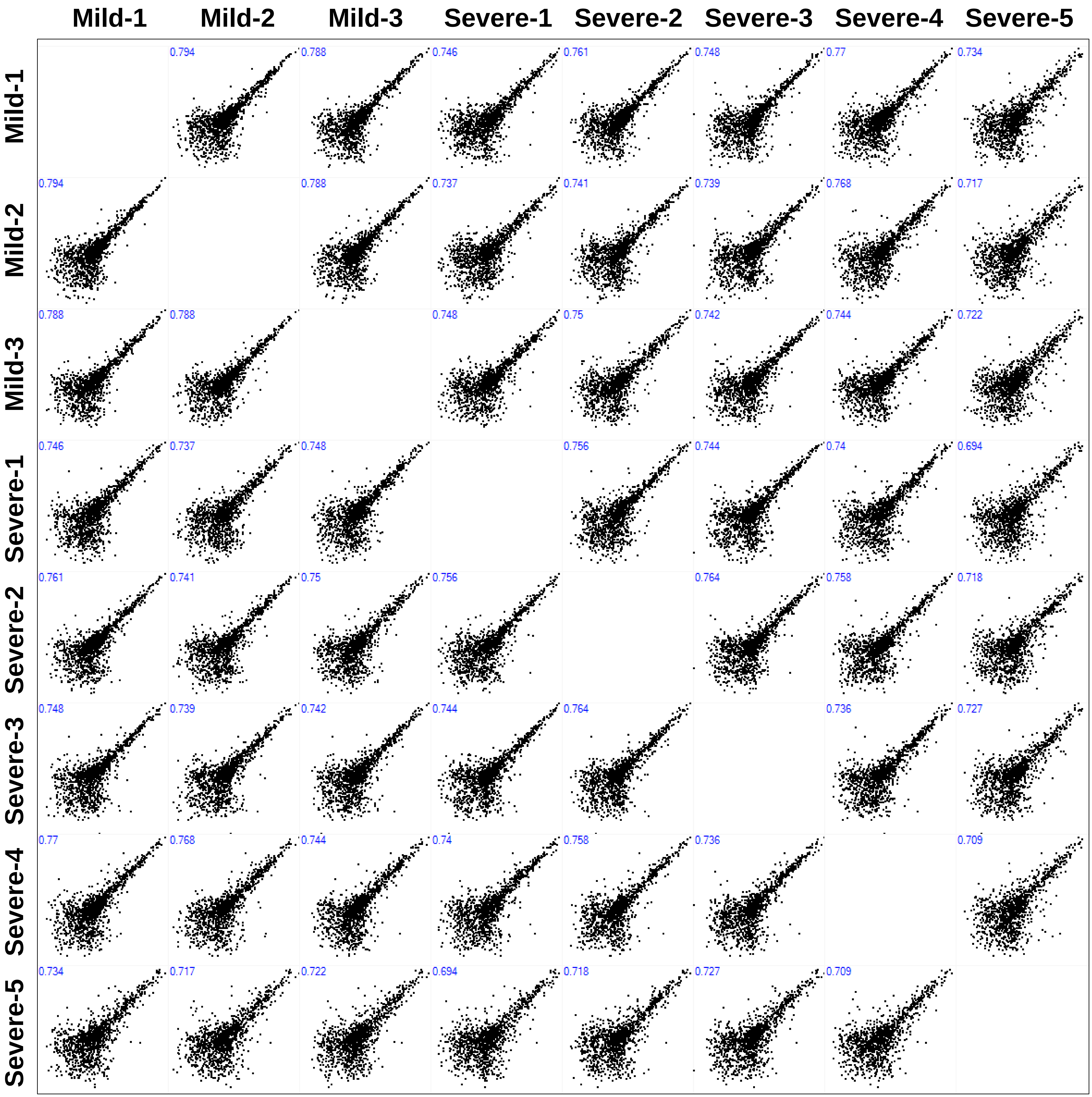


**Figure S1. Correlation plot in plasma experiment**

The scatter plot showed the correlation between the samples. Pearson's correlation coefficient value were calculated for each same pair and displayed.

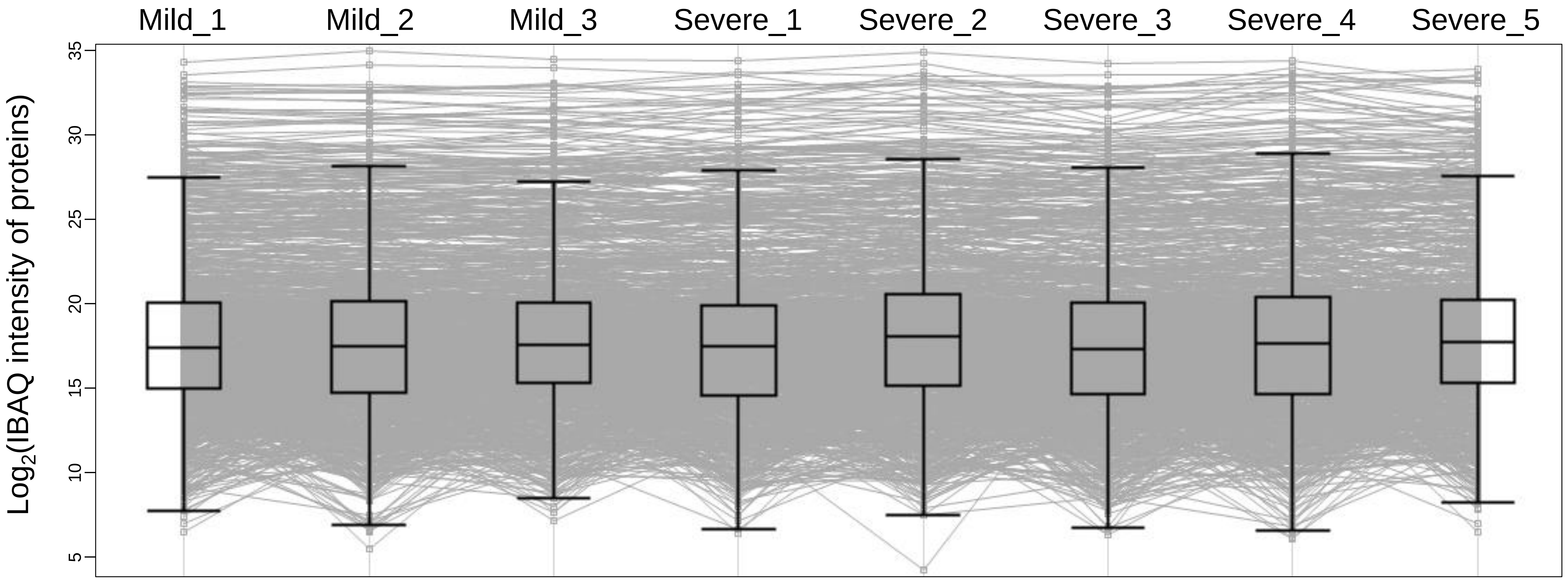


**Figure S2. Profile Plot of all proteins across the plasma samples**
The log-transformed abundance of all proteins were plotted across the plasma samples. The five-number summary of box-and-whisker is the 5%, first quartile, median, third quartile, and 95%.

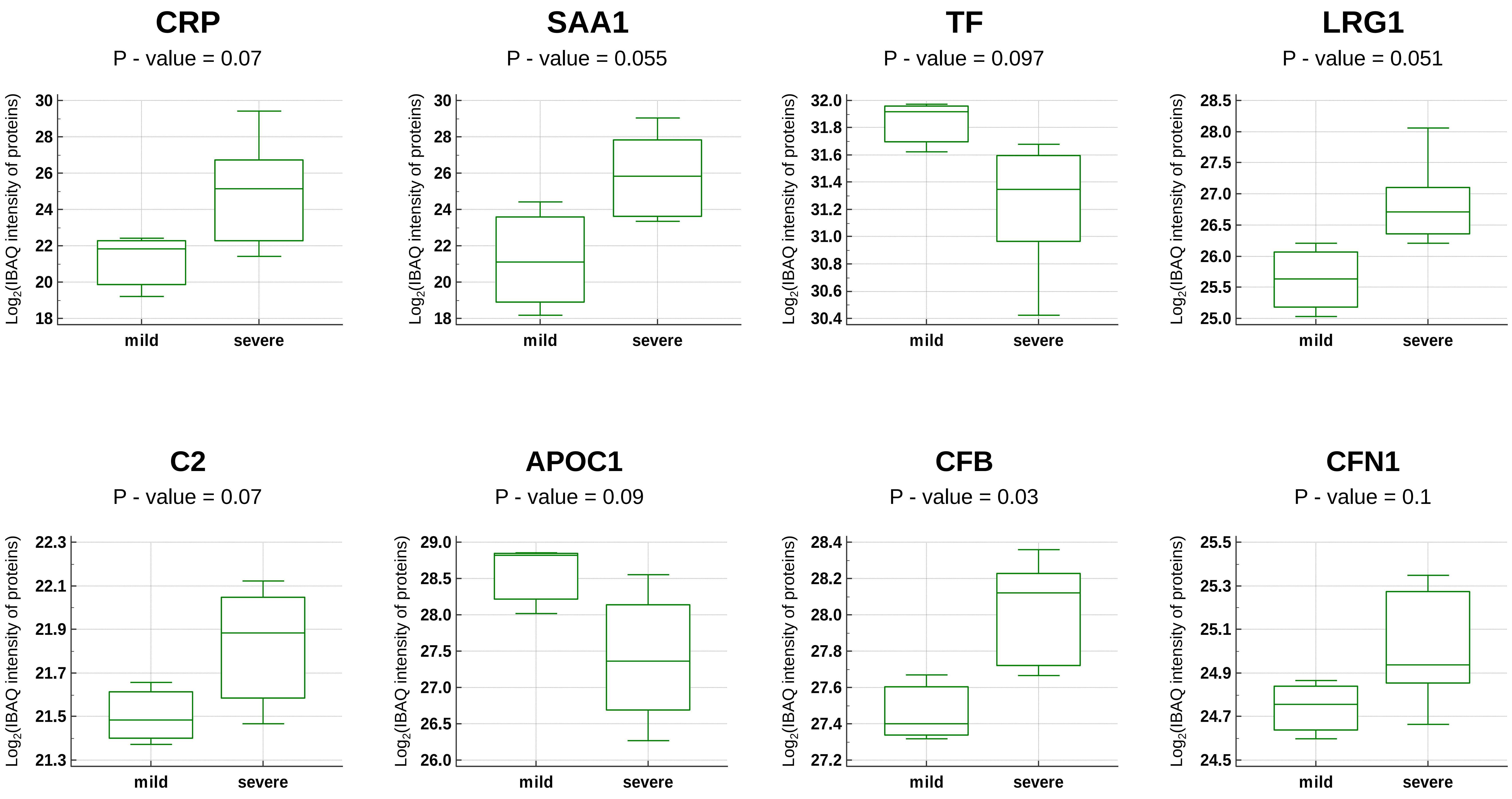


**Figure S3. The expression summary for the proteins reported as significant in previous papers**

The log-transformed abundance of the proteins which were not significant (0.05 < P value < 0.1 or Fold-change < 1.5) in our data but significant commonly in other datasets (Figure 3B) were plotted against the severity groups. The five-number summary of box-and-whisker is the minimum, first quartile, median, third quartile, and the maximum.

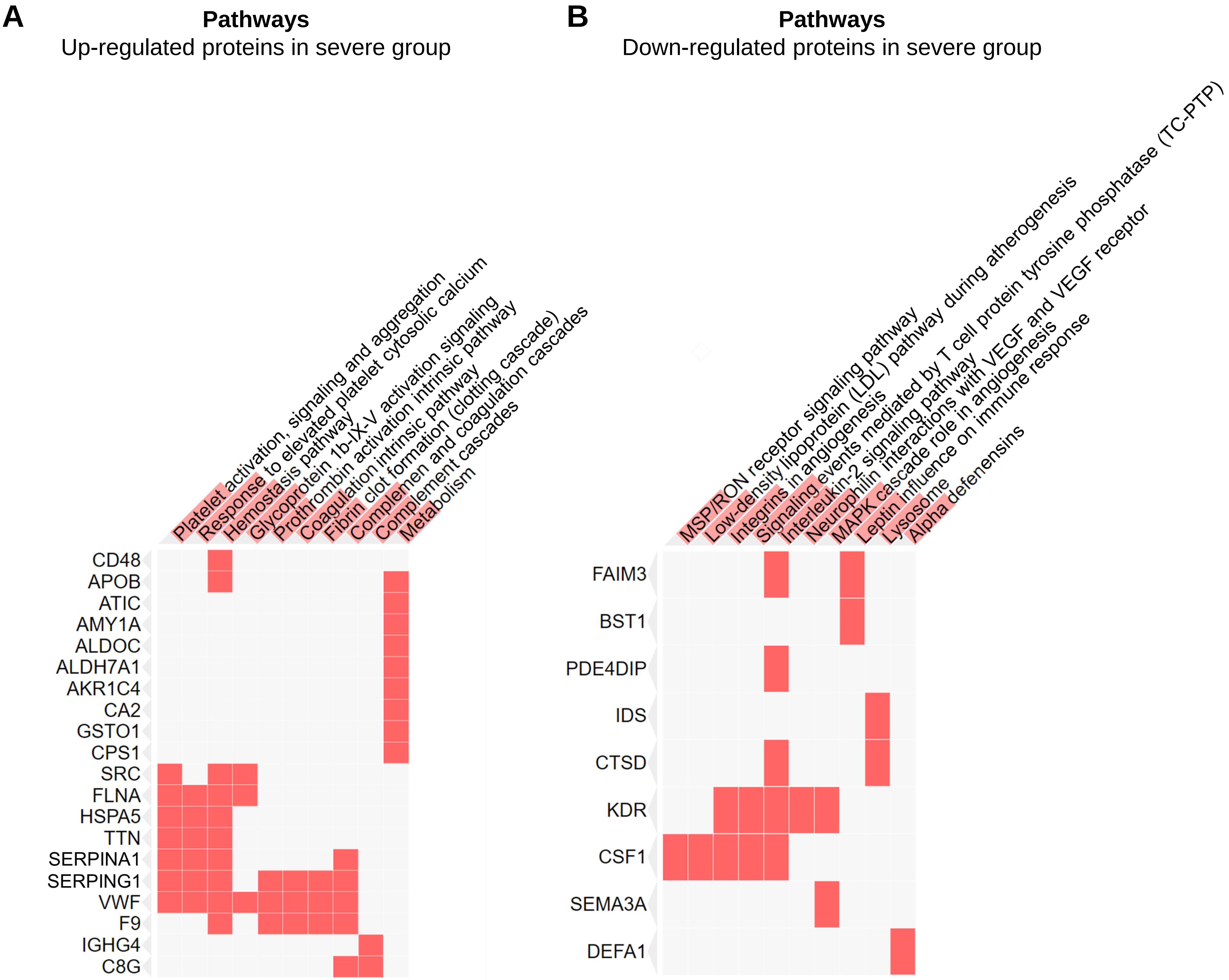


**Figure S4. Pathway enrichment using up- or down-regulated DEPs**

A, B. Pathway enrichment analysis was performed using the up- or down-regulated DEPs, respectively. If the proteins in left side were associated in corresponding pathway, they were filled with red. The level of significance of each pathway is displayed as the length of red bar overlapped in the pathway name cell.
